# Mobile retroelements induced by hypomethylating agents are restricted to transpose in myeloid malignancies

**DOI:** 10.1101/2023.07.30.551176

**Authors:** Šárka Pavlová, Marcela Krzyžánková, Anastasiya Volakhava, Anastasia Smirnova, Tatiana Grigoreva, Zuzana Jašková, Hana Synáčková, Dennis Wahl, Michaela Bohúnová, Libor Červinek, Šárka Pospíšilová, Ilgar Mamedov, Karla Plevová

## Abstract

Retroelements (RE) present in the human genome are silenced via multiple mechanisms, including DNA methylation, to prevent their potentially mutagenic effect. RE activity, demonstrated by their expression and somatic retrotransposition events, is deregulated in multiple tumor types but not in leukemia. We hypothesized that treatment with hypomethylating agents (HMA), commonly used in myelodysplastic syndromes and acute myeloid leukemia, could lead to increased RE activity and somatic retrotranspositions, and contribute to disease progression. We induced expression of ORF1p protein encoded by long interspersed nuclear element-1 (L1) after 72h treatment with HMA in DAMI and HL-60 cell lines. ORF1p was predominantly localized in the cytoplasm, as evidenced by fluorescent microscopy of the DAMI cell line. To study whether long-term HMA therapy may induce somatic retrotranspositions, we (i) treated both cell lines for four weeks, (ii) analyzed a cohort of 17 MDS patients before and on treatment with HMA. Using a previously established sensitive NGS-based method, no RE events were identified. To conclude, we show that although HMA induces the expression of L1-encoded proteins in tumor myeloid cell lines, *de novo* somatic retrotransposition events do not arise during the long-term treatment of MDS patients and myeloid cell lines with these agents.

## Introduction

Long interspersed nuclear elements (LINEs), a non-LTR subfamily of retrotransposons, occupy approximately 17% of the human genome (1). Out of around 500 000 copies, the majority are inactive due to genomic rearrangements, point mutations, and 5ʹ-truncation to prevent their harmful impact on the genome stability. The ongoing transposition is primarily caused by type 1 (LINE-1; also known as L1). Only around 100 copies of L1 elements per genome are responsible for the majority of retrotransposon activity (2, 3).

The L1 transposable element comprises two open reading frames, ORF1, encoding an RNA chaperone, and ORF2, encoding an enzyme with single-strand endonuclease and reverse transcriptase activities. After L1 transcription, the polyadenylated bicistronic L1 mRNAs are transported to the cytoplasm and translated into ORF1p and ORF2p proteins. Multiple ORF1p trimers and one or two ORF2p molecules bind the mRNA in cis to form the L1 ribonucleoprotein (L1 RNP). The strong predominance of ORF1p proteins (∼30x) explains their much easier detection compared to ORF2p (4). Indeed, ORF2p is nearly undetectable in primary tumors (5). L1 RNP penetrates the nucleus during mitosis (6). Subsequently, target-primed reverse transcription occurs during the S-phase, initiated by the endonuclease nicking genomic DNA at A- and T-rich sites, followed by annealing of RNA poly(A) tail and its reverse transcription (7, 8). Using host proteins involved in DNA repair and replication, the single-stranded DNA gap at the L1 integration site is recognized, followed by the integration and ligation of a newly synthesized L1 copy into the DNA, being flanked by the target site duplications.

The process of retrotransposition is suppressed by multiple cellular mechanisms, including DNA methylation, histone acetylation, piwi RNA complexes, and p53 machinery, with a frequency of a new retrotransposition event in the human population being between 1/20 and 1/200 births (9, 10). As such, many L1 elements are present in individual genomes but absent in the haploid human genome reference. Contrary to normal cells, the expression levels of transposable elements and corresponding proteins, including ORF1p, are increased in tumor cells (11, 12). Consequently, L1 retrotranspositions have been documented in multiple cancer types (13–15). Active TEs are considered highly mutagenic and are commonly linked to the multiple steps of cancer development and progression (14, 16–18). Moreover, L1 hypomethylation has been revealed in multiple cancer types, including lung cancer (19), colorectal cancer (20), breast cancer (21), prostate cancer (22), hepatocellular carcinoma (23), ovarian cancers (24) and esophageal squamous cell carcinoma (25), and was typically associated with poor clinical outcomes, likely due to resulting genome instability and presumed TE activation.

While high levels of somatic retrotranspositions have been shown mainly in solid tumors, in hematological malignancies, the levels are considered to be generally low (15). In line, we did not identify new tumor-specific RE insertions in childhood and adult acute leukemia samples using a sensitive NGS approach allowing for the detection of Alu and L1 insertions in 1% of cells (26).

Hypomethylating agents (HMA) 5-Azacytidine (Aza) and 5-Aza-2ʹ-deoxycytidine (decitabine, Aza-dC), either alone or in combination with other drugs, have become standard therapy for patients with high-risk myelodysplastic syndromes (MDS) (27, 28) and acute myeloid leukemia (AML) (28). Despite the unequivocal benefit for overall survival as well as improved quality of life, some patients do not respond to these drugs, and the majority of responding patients relapse. Since their use leads to hypomethylation of silenced regions in the human genome, including L1, we studied if HMA affect the activity of L1 retrotransposons in tumor myeloid cell lines and if *de novo* L1 retrotransposition can be detected in MDS patients treated with and progressing after Aza.

## Materials and methods

### Cell lines and patient samples

A panel of tumor cell lines (Table S1) was used for the initial screening of ORF1 expression following cultivation instructions provided by the supplying collection (DSMZ or ATCC). For further experiments, human myeloid cell lines DAMI and HL60 were selected. The breast carcinoma MCF-7 cell line was used as a positive control for ORF1 expression. HEK293T/17 cell line was used for transient transfections.

Thirty-seven serial bone-marrow samples from 17 MDS patients treated with 5-azacytidine (Aza) were taken after written informed consent approved by the Ethical Committee of the University Hospital Brno was available following the Declaration of Helsinki. The samples were taken before treatment initiation, after several lines of Aza treatment, and/or in relapse; if a relapse sample was unavailable, the closest available sample before progression was used (Table S2). DNA was isolated from whole bone marrow leukocytes after erythrolysis.

### Treatment of cell lines with 5-Aza-2ʹ-deoxycytidine

To induce ORF1p and ORF2p expression, a panel of tumor cell lines (Table S1) was treated with 5-Aza-2ʹ-deoxycytidine (Aza-dC; Sigma-Aldrich) for 72 hours (0.5, 1, 2 and 5 μM). Briefly, 2.5 × 10^5^ cells/ml were seeded in 6-well plates (TPP) in 5 ml of cell culture medium (Table S1) + 10% fetal bovine serum (FBS). On the following day, medium was replaced with fresh medium containing Aza-dC (0.5, 1, 2, and 5 μM) and cultured for 48 hours, then the treatment was repeated and followed by cultivation for additional 24 hours. As a control, the cell lines were processed in the same way without adding Aza-dC.

For fluorescent microscopy and flow cytometry, DAMI and HL-60 were seeded in 6-well plates (TPP) with or without sterile microscopy coverslips in density 3.0 × 10^6^ cells/ml in 3 ml of DMEM + 10% ultra-low IgG FBS (PAN-biotech). On the following day, the cells were treated with 2 μM and 5 μM Aza-dC with media change after 48 hours, as described above.

Long-term treatment of DAMI and HL-60 was performed with 0.5 and 2 µM of Aza-dC with untreated cells as controls. Cells were harvested at days 0, 3, 7, 14 and 28.

### Immunoblotting

Cells were washed with PBS and lysed in ice-cold NP-40 buffer (Thermo Fisher Scientific) with protease and phosphatase inhibitors (1:100; P8340 and P0044, Sigma Aldrich) for 30 minutes. Protein concentrations were determined by Bicinchoninic Acid Kit for Protein Determination (BCA1-1KT, Sigma Aldrich) or the Bradford Protein Assay (BioRad). The protein lysates were run on 10% sodium dodecyl sulfate-polyacrylamide gels (SDS-PAGE) and transferred to a nitrocellulose membrane (Bio-Rad) using a wet tank blotting system. The membranes were blocked with 5% non-fat milk in TBS buffer containing 0.1% Tween on a rocking shaker for 2 hours. The membranes were washed in TBS-T alone and incubated in the appropriate primary antibody overnight on a rocking shaker at 4 °C. The following day, membranes were washed and incubated in the appropriate secondary antibodies at RT for 1 hour. For the list of antibodies, see Table S3. The proteins were detected using a chemiluminescence system (Clarity Western ECL Substrate, 1705061, Bio-Rad) and imaged using a UVITEC Documentation System (Uvitec Cambridge, Mini HD9).

### Intracellular detection of ORF1p and ORF2p proteins using fluorescence microscopy and flow cytometry

For fluorescent microscopy, the sterilized coverslips (13 mm/0.17 mm Menzel) coated with 0.01% poly-L-lysine (Sigma-Aldrich) were seeded with the targeted cell line at a volume of 200 µl per well.

Cultured cells were fixed with 4% formaldehyde (10 min; F8775, Sigma-Aldrich) and blocked with 3% IgG-free BSA blocking buffer (001-000-161, Jackson ImmunoResearch) with the addition of 0.25% Triton X-100 (SIALX100, Sigma-Aldrich) for 1 hour. The cells were incubated in the corresponding primary antibody for 1 hour and the secondary antibody for 30 min (Table S3). Nuclear DNA was stained with Hoechst (H1399, ThermoFisher) for 2 min. All procedures were performed at room temperature. Finally, individual stained coverslips with cells were carefully transferred with tweezers to slides with prepared mounting medium (S3023, DAKO). Fluorescence detection of ORF1p and ORF2p was performed the day after the mounting medium dried using a ZEISS 700LSM confocal microscope with plan-apochromat 40x oil objective lens. Individual images were detected using two filters with 405 and 488 nm wavelengths and 1 AU pinhole and assessed in ZEN 2009 software.

Intracellular staining for flow cytometry using FACSVerse (BD Biosciences) was performed analogically to fluorescent microscopy staining without coverslips.

### Preparation of ORF1 and ORF2 Plasmids and transient transfection

*E.coli* DH5alpha with pBudORF1 (Addgene) and pBudORF2 (Addgene) were cultured according to the manufacturer’s protocol. Plasmid DNA was isolated using EndoFree Plasmid Maxi Kit (Qiagen) according to the manufacturer’s protocol.

Transient transfection for fluorescence microscopy was performed directly on coverslips coated with poly-L-lysine (see above) placed in 24-well plates using the transfection reagent Polyethylenimine (PEI; Polysciences) according to the manufacturer’s protocol. Briefly, the day before transfection, HEK293T/17 cells were seeded in individual wells of a 24-well plate with coverslips at a cell density of 0.5 × 10^6^/ml with 1 ml of DMEM/F12 culture medium (PAN-biotech) and 10% ultra-low IgG FBS (PAN-biotech). On the day of transfection, the culture medium was replaced with DMEM/F12:H2O transfection medium in a 1:1 ratio with the addition of 300 µl serum-free 2mM Glutamine (PAN-biotech). The premix was added to the individual wells with prepared transfection media after being left in the box at RT for half an hour at a DNA:PEI ratio (3:9 µg) of 150 µl. The transfected cells were incubated at 37 °C and 5% CO_2_. After 4 hours, the transfection medium was replaced with fresh 1ml DMEM/F12 medium + 10% ultra-low IgG FBS and plates were incubated for additional 72 hours. Fluorescence staining of the slide was carried out after PBS wash, as described above.

Transient transfection for the detection of ORF proteins by flow cytometry and western blots was performed in 6-well plates without coverslips with the identical transfection procedure, and the volumes increased accordingly, i.e., 1.5 x 10^6^ in 3 ml of HEK293T/17 cells were used, and the volumes of culture medium and transfection medium were 1 ml and 500 μl, respectively. After three days of culturing in 3 ml of fresh medium, cells were washed out with PBS supplemented with 15 mM EDTA and transferred into 1.5 ml Eppendorf tubes.

### Detection of DNA retrotransposition insertions in cancer cell lines and primary patient cells

A next-generation sequencing method for detecting new L1 insertions of the transpositionally-active subfamily L1HS was used according to our previous reports (26, 29). Briefly, gDNA was digested by a mixture of selected endonucleases TaqI and FspBI to generate fragments that consisted of a 3’ part of L1 retroelement and its adjacent genomic region (i.e., flank). In the next step, the fragmented DNA was ligated to a stem-loop adapter containing unique molecular identifiers (UMI) that were used to quantify the number of cells bearing each insertion after sequencing. Next, a primer specific to transpositionally active L1 subfamily L1HS and a primer corresponding to the ligated adaptor were used to selectively amplify genomic flanks adjacent to the 3’ end of L1. A product of the first PCR was used in the second semi-nested PCR. Finally, an indexing PCR was carried out to introduce the sample barcodes and oligonucleotides necessary for Illumina sequencing on the NextSeq 550 machine (paired-end, 150+150). We used a custom computational pipeline (30) to map all the sequenced insertions’ flanks to the human reference genome and remove various artifacts generated during library preparation and sequencing. The coordinates of the insertions in the serial samples were matched to identify insertions related to clonal propagation (i.e., compared to the corresponding pre-treatment sample). Following our previous report (26), candidate somatic L1 insertions were validated by an independent locus-specific PCR with initial gDNA as a template, and Encyclo Polymerase Mix (Evrogen) and/or Q5 High-Fidelity 2X Master Mix (New England Biolabs).

For detailed protocols, see *Supplementary Methods*.

## Results

### 5-Aza-2ʹ-deoxycytidine treatment induces ORF1p expression in tumor myeloid cell lines

First, we performed initial Western blot screening to reveal which cell lines express ORF1p (Table S1). While ORF1p was clearly detected in three carcinoma cell lines (MCF-7, SW-48, H1299), it was absent in all seven lymphoid and all four myeloid leukemia cell lines. Since DNA methylation is a key transposon silencing mechanism, we attempted to induce ORF1p expression using Aza-dC. Out of the tested cell lines, we were able to induce ORF1p expression in two myeloid cell lines, DAMI and HL-60 (Figure 1A), while for the remaining two cell lines derived from myeloid disorders used Azd-dC dosage led to a prominent decrease in cell viability (data not shown). Thus, DAMI and HL-60 cell lines were used for further experiments. To validate our results with an independent method, we implemented a protocol for intracellular ORF1p staining and analyzed treated and untreated DAMI and HL-60 cells with flow cytometry. We confirmed the ORF1p induction in both tested cell lines (Figure 1B). Similarly to the ORF1p levels detected by the Western blot analysis in bulk samples, we did not observe a dose-dependent effect when increasing Aza-dC concentrations from 1 to 5 μM. Next, to explore the intracellular localization of ORF1p in myeloid cell lines, we stained the cells intracellularly on microscope coverslips for visualization using fluorescent microscopy. As positive controls, we used MCF-7 with a high endogenous expression (Figure S1) and HEK293T/17 cells transfected with pBudORF1 expression plasmid (Figure S2A). The optimized procedure was used in DAMI cells that could adhere to the surface (unlike HL-60). Again, we confirmed ORF1p induction in Aza-dC-treated cells (Figure 2). ORF1p signals predominated in the cytoplasm, with scarce presence in the nuclei.

**Figure 1.**
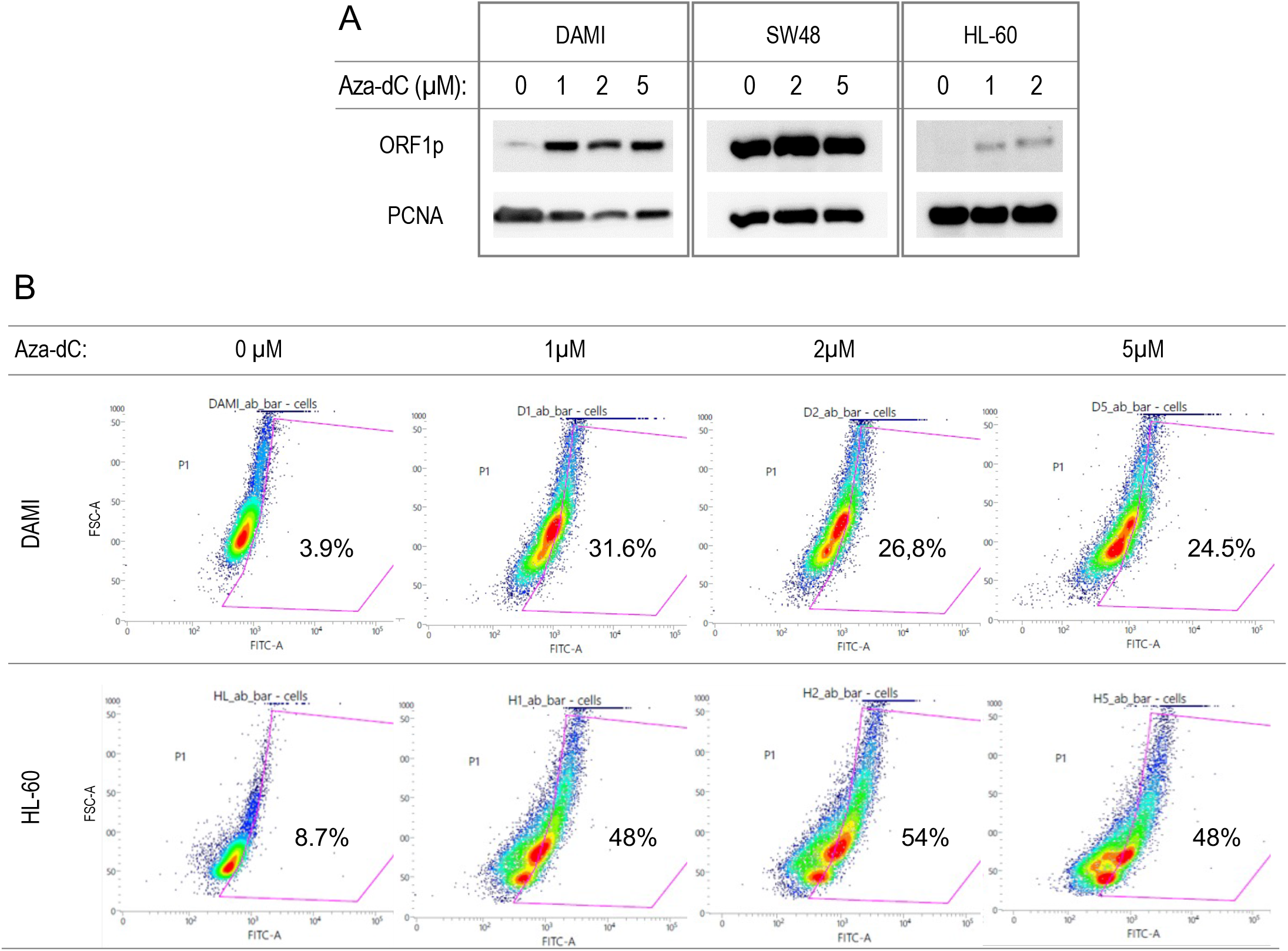
Western blot (A) and flow cytometry (B) measurements of ORF1p after Aza-dC treatment. The Western blot analysis was performed on a panel of tumor cell lines (Table S1). Here only DAMI and HL-60 (myeloid cell lines) and SW48 (colorectal carcinoma cell line) are shown to demonstrate the ORF1p induction in the myeloid cell lines, while in SW48, the ORF1p expression was present already in the baseline sample without Aza-dC treatment. The ORF1p induction in DAMI and HL-60 was further confirmed using flow cytometry. Aza-dC concentrations applied for the Western blot analysis – DAMI: 0, 1, 2, 5 µM; SW48: 0, 2, 5 µM; HL-60: 0, 1, 2 µM; and for flow cytometry – both DAMI and HL-60: 0, 1, 2, 5 µM. Primary/secondary antibodies – Western blot: Anti-LINE-1 ORF1p (CellSignalling); flow cytometry: Anti-LINE-1 ORF1p (Abcam) + Goat anti Rabbit IgG AF488 (Abcam).

**Figure 2.**
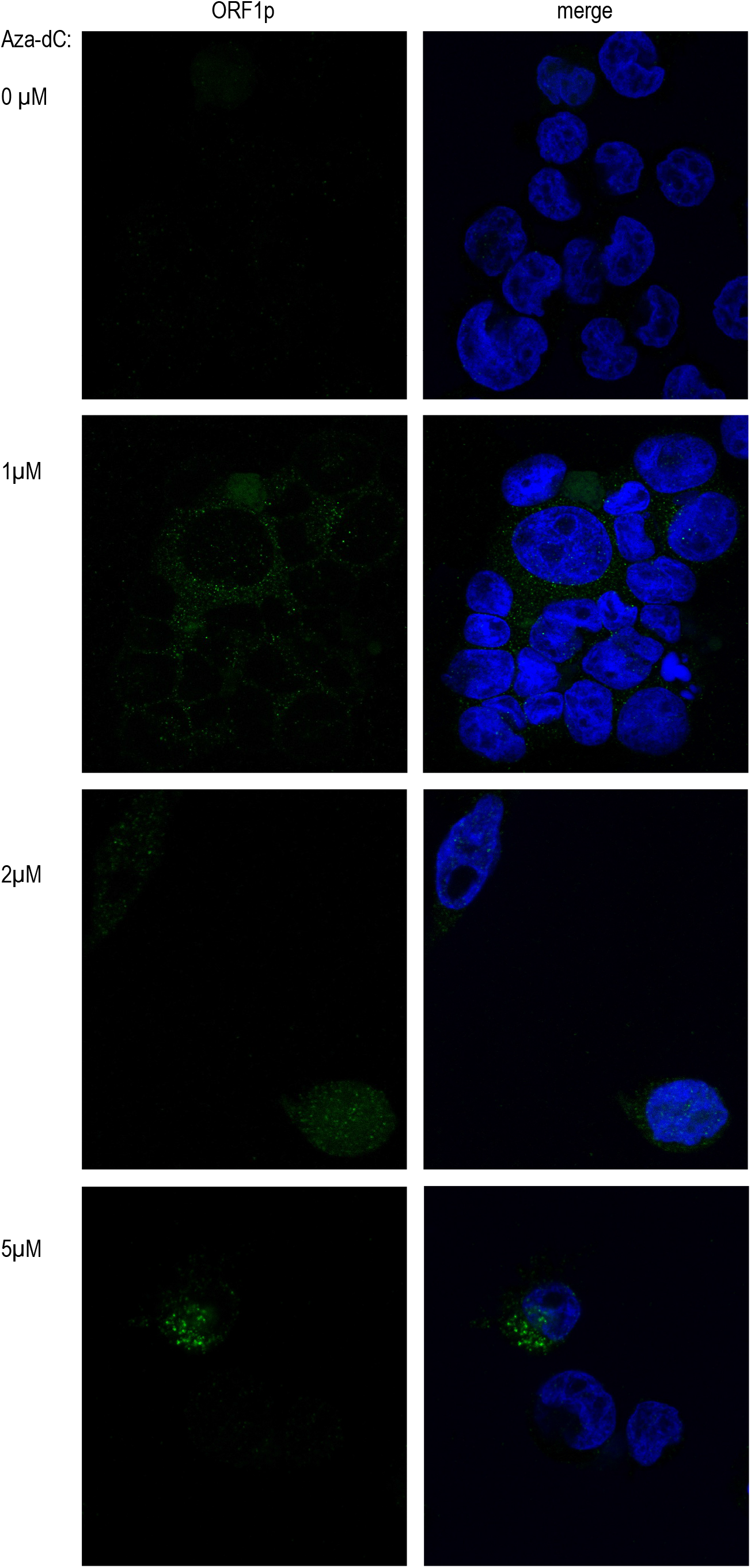
Confocal microscopy detection of ORF1p induction after Aza-dC treatment in DAMI cell line. Left panel: The immunofluorescence signal of ORF1p (green). Right panel: The immunofluorescence signal of ORF1p (green) combined with the nuclei signal (DAPI, blue). Primary/secondary antibodies: Anti-LINE-1 ORF1 (Abcam) + Goat anti Rabbit IgG AF488 (Abcam). Magnification: 40x.

Further, we also aimed to study the expression of ORF2p, a 150 kDa protein with much lower cellular abundance. For this purpose, we treated the DAMI cell line with Aza-dC and analyzed it using flow cytometry. Although we observed a protein induction after Aza-dC treatment (Figure 3), we were unable to confirm the induction with the fluorescence microscopy despite successfully detecting ORF2p in HEK293T/17 cells transiently transfected with pBudORF2 expression plasmid (Figure S2B), likely due to different sensitivity of the two methods.

**Figure 3.**
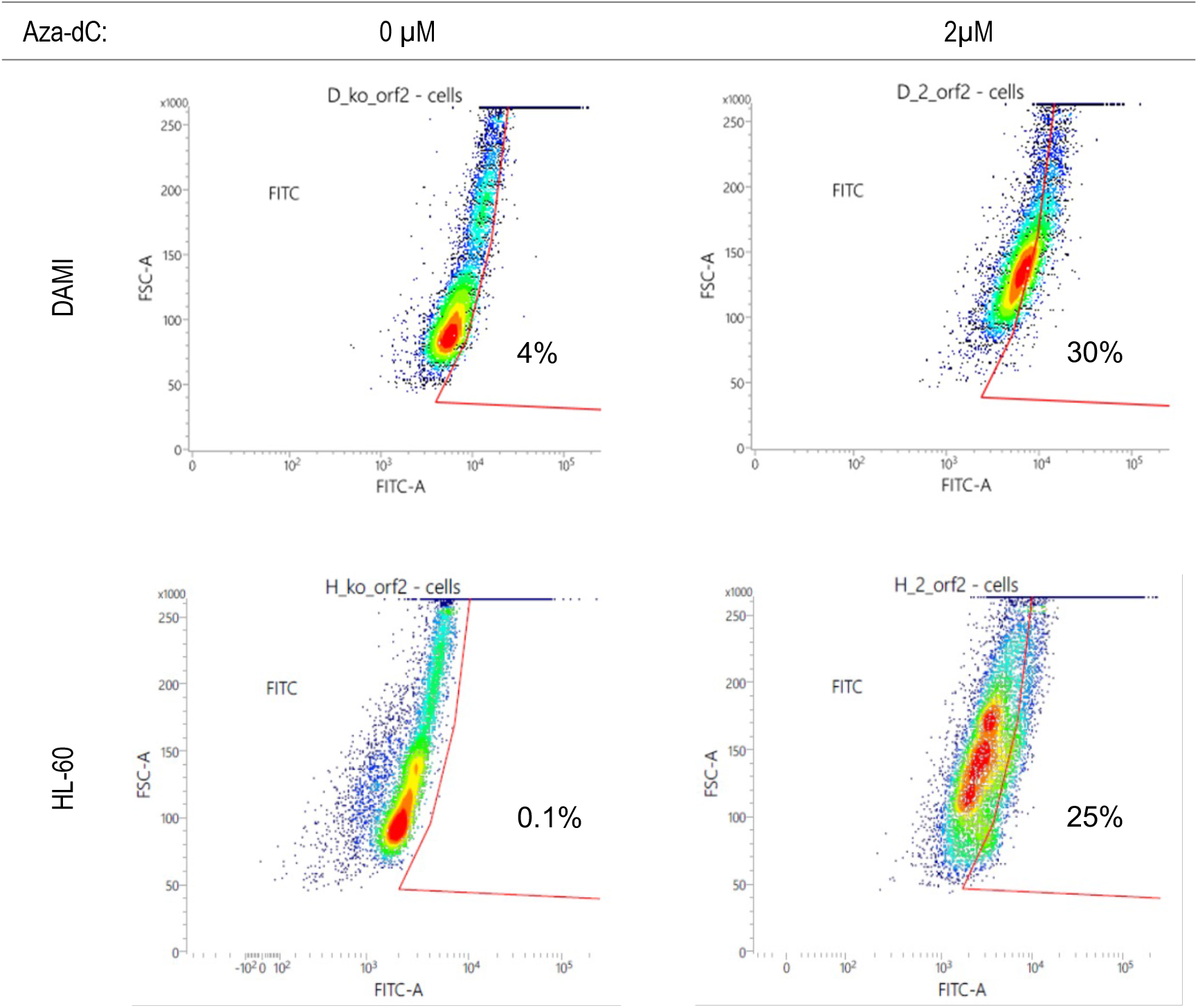
Flow cytometric analysis of ORF2p induction after the treatment with 2 µM Aza-dC in DAMI and HL-60 cell lines. The experiment was repeated in independent biological triplicate or duplicate for DAMI and HL-60, respectively, with similar results. Primary/secondary antibodies: Anti-LINE-1 ORF2 (Rockland) + Goat anti Chicken IgY AF488 (Invitrogen).

### Long-term treatment of tumor myeloid cell lines with 5-Aza-2ʹ-deoxycytidine does not increase the retrotransposition rate

Based on our findings, showing that HMA may induce the expression of proteins encoded by L1 retrotransposon, we studied whether the long-term exposition of myeloid tumor cells to HMA also increases the retrotransposition rates. To mimic *in vitro* the therapy of hematooncology patients, we treated DAMI and HL-60 cell lines with 0.5 and 2 μM Aza-dC for a period of four weeks. On days 3, 7, 14, and 28, we collected DNA, protein and cells.

First, we assessed ORF1p induction during Aza treatment using Western blot and flow cytometry and observed the level of induction for 2 μM Aza comparable with our initial experiments (data not shown). This result prompted us to proceed with detecting novel insertions using NGS. We applied a high-sensitive NGS-based protocol enabling us to detect L1 insertions down to 1% of cells (26). Candidate *de novo* L1 insertions for each time point (3, 7, 14, and 28 days after Aza-dC addition) were determined by comparing their genomic coordinates to the list of L1 insertion sites identified in the same cell line before treatment (day 0). We additionally filtered out the insertions whose coordinates matched the insertions found in other treatment conditions. Such insertions comprise false positives as the probability of an independent insertion into the same genomic locus in different cells is infinitely low. Following our pipeline, manual filtering against the genomic sequence, and excluding regions containing known population L1 copies, we identified 358 candidate insertions. Based on read quality (R1 and R2 matching) and number of reads (>10) and UMIs (≥2), we selected 14 candidate insertions (8 in DAMI and 6 in HL-60), for which we designed locus-specific primers for PCR validation (Table S4, Figures S3A-B). None of the candidate insertions was confirmed.

### Retrotransposition is uncommon in MDS patients treated with HMA

Still, we wanted to find out whether we could identify any retrotranspositions in leukocyte DNA of MDS patients treated with Aza and whether the clonal expansion of cells carrying *de novo* L1 insertions may contribute to disease relapse. The rationale behind that was that they were exposed to the hypomethylation effect for a much longer period than the cell lines in four-week-lasting cultivation and the tumor cells eventually escape from the cell death caused by Aza. For this purpose, we again used our high-sensitive NGS-based protocol to explore 37 serial bone-marrow samples from 17 MDS patients treated with Aza. The cohort consisted of cases with high-risk disease, with five patients bearing *TP53* defects and nine progressing to AML. The samples were taken before treatment initiation, several months on treatment and/or in relapse (for details about samplings, see Table S2). The patients received a median of 11 Aza cycles (ranging from 3 to 46) between the baseline and last samples. L1 insertions of each follow-up sample were compared to the respective baseline sample of the same patient.

Following the same manual filtering as for treated tumor cell lines, we identified three candidate insertions in the genomes of three MDS patients undergoing therapy, each bearing a single candidate insertion, that were subjected to validation using locus-specific PCR (Figure 3, Table S5). Out of these candidate insertions, we were able to validate none of them, suggesting that L1 retrotransposition activity in primary patient samples is low.

## Discussion

Epigenetic silencing of transposable elements is one of the key mechanisms of cellular defense against their potentially mutagenic effect. As REs become active after demethylation of their promoters (12, 23), we aimed to explore whether they become active in MDS treated with HMA and whether *de novo* insertions contribute to disease progression, at least in some patients.

MDS is a heterogeneous disease comprised of hematopoietic stem cell disorders leading to ineffective hematopoiesis, demonstrated by blood cytopenias and progression to acute myeloid leukemia in approximately one third of patients (31, 32). Besides cytogenetic changes and gene mutations, epigenetic changes contribute to disease pathophysiology and progression. Widespread hypermethylation was correlated to impaired overall survival and progression to AML (33, 34). Although HMA are highly efficient in MDS, the mechanism of their action is not fully understood. It includes activation of tumor suppressor genes commonly inactivated in MDS, incorporation of modified cytidine derivates into RNA and DNA, and activation of tumor antigens and retroelements recognized by the immune system. (Reviewed by (35) and (36)).

Indeed, we demonstrated with several methods that ORF1p levels increase after Aza-dC treatment of tumor myeloid cell lines DAMI and HL-60, pointing to retroelement activation. Fluorescent microscopy showed that the majority of ORF1p induced by Aza-dC in DAMI is localized in the cytoplasm and scarcely in the nucleus. We were unable to localize ORF2p after treatment and could only show its expression and localization in the ectopic expression model. This is in line with other studies (5) and can be partially explained by the predominance of ORF1p proteins in L1 RNP (4). As the Aza-dC caused cell death in other myeloid cell lines tested, we could not study ORF1p and ORF2p induction and localization in other models.

Despite the apparently increased level of L1 proteins, we did not observe *de novo* retrotranspositions in cell lines and patients treated with HMA. This may be explained by the absence of a strong selective advantage that would lead to clonal expansion of affected cells. However, as we expected that novel L1 insertions could be present in minor subpopulations and the whole genome sequencing methods would have limited sensitivity to detect them, we applied a targeted-NGS-based approach with the ability to capture retrotransposition events in 1% of cells (26). This approach was previously used to demonstrate somatic retrotranspositions in brain tissue (37) and colorectal cancer (29) and the absence of RE activity in leukemia (26). Thus, we assume that the sensitivity of the NGS method was not the reason for the negative result.

The alternative explanation for the absence of L1 retrotransposition can be that myeloid tumor cells are negatively selected for L1 expression and/or the retrotransposition process. Indeed, a comprehensive study showed that decreased L1HS expression is associated with short overall survival in AML, and it has been suggested that L1 silencing is required for oncogene-induced transformation and propagation of AML-initiating cells (38). In a model system, activation of endogenous L1s resulted in increased L1 retrotransposition and impaired leukemia cell growth *in vitro* and in xenotransplants. In line, a model proposed by (39) suggests that the impact of TEs activity evolves during myeloid transformation. While the increased activity of TEs and piRNAs (40) induces an immune response and leads to the elimination of leukemic cells in early-stage MDS, the leukemic cells finally escape the control of the immune system via suppression of TE/piRNA expression, resulting in the progression to high-risk MDS, accumulation of leukemic blasts, and AML.

The somatic retrotranspositions are absent not only in AML and high-risk MDS but also in acute lymphoblastoid leukemia, as we showed previously (26). Of note, in the report, we also analyzed a set of childhood AML that are biologically distinct from AML in adults and the elderly, and we did not detect somatic retrotranspositions either (26). Whether the intolerance of RE activity is a feature of hematopoietic stem and early progenitor cells or is inherent to the hematopoietic lineage as a whole is currently unknown. It would be of interest to explore RE activity, e.g., in mature B-cell neoplasms that are not directly related to a hematopoietic stem cell.

Significantly increased levels of ORF1p proteins and L1Hs transcripts were detected in TP53-mutated Wilms tumor and colorectal carcinoma samples, respectively (41). It has been suggested that keeping genome integrity via preventing RE events is an ancient function of tumor suppressor p53 (42). *TP53*-mutated myeloid neoplasms have recently been recognized as an aggressive entity (43). To see if loss of p53 function enables RE events in MDS, we included five patients with *TP53* mutations in the studied cohort. The lack of RE events in these patients and the *TP53*-deficient cell lines HL-60 and DAMI suggests that intolerance of RE events in leukemic cells is p53-independent, although it is to be confirmed by future studies. Overall, we show that although HMA induce the expression of L1-encoded proteins in tumor myeloid cell lines, *de novo* somatic retrotransposition events do not arise during the treatment of MDS patients with these agents. Thus, while somatic retrotranspositions occur in carcinomas, we show they are uncommon in myeloid cells.

## Author contributions

SPav, IM, and KP conceived the study, designed and supervised experiments, and interpreted data; MK and IM developed methods; MK, AV, TG, ZJ, HS, and DW performed experiments and analyzed data; AS carried out the bioinformatic analyses; MB collected clinical and laboratory diagnostic information; LC provided samples and clinical expertise; SPav, IM, and KP drafted the manuscript; SPosp, IM and KP obtained funding; all authors reviewed and approved the manuscript.

## Data Availability Statement

Raw sequencing data are available on the Sequence Read Archive (SRA) under the accession no. SUB13719421.

