## Supplementary Figures 1-3 for "Mobile retroelements induced by hypomethylating agents are restricted to transpose in myeloid malignancies"

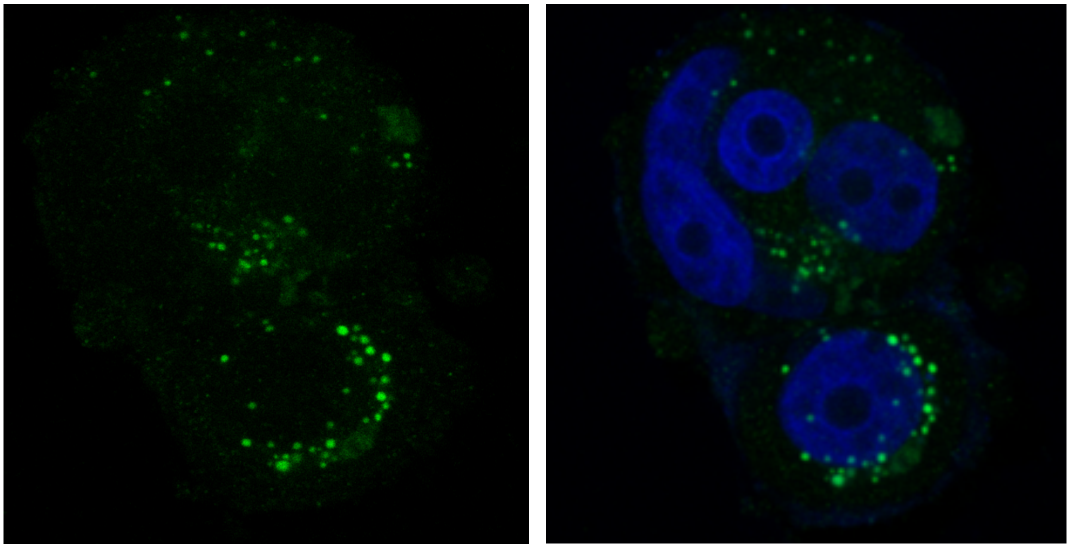

**Figure S1**. Confocal microscopy detection of ORF1p in MCF-7 cell line used as a positive control. Left: The immunofluorescence signal of ORF1p (green). Right: The immunofluorescence signal of ORF1p (green) combined with the nuclei signal (DAPI, blue). Primary/secondary antibodies: Anti-LINE-1 ORF1 (Abcam) + Goat anti Rabbit IgG AF488 (Abcam). Magnification: 40x.

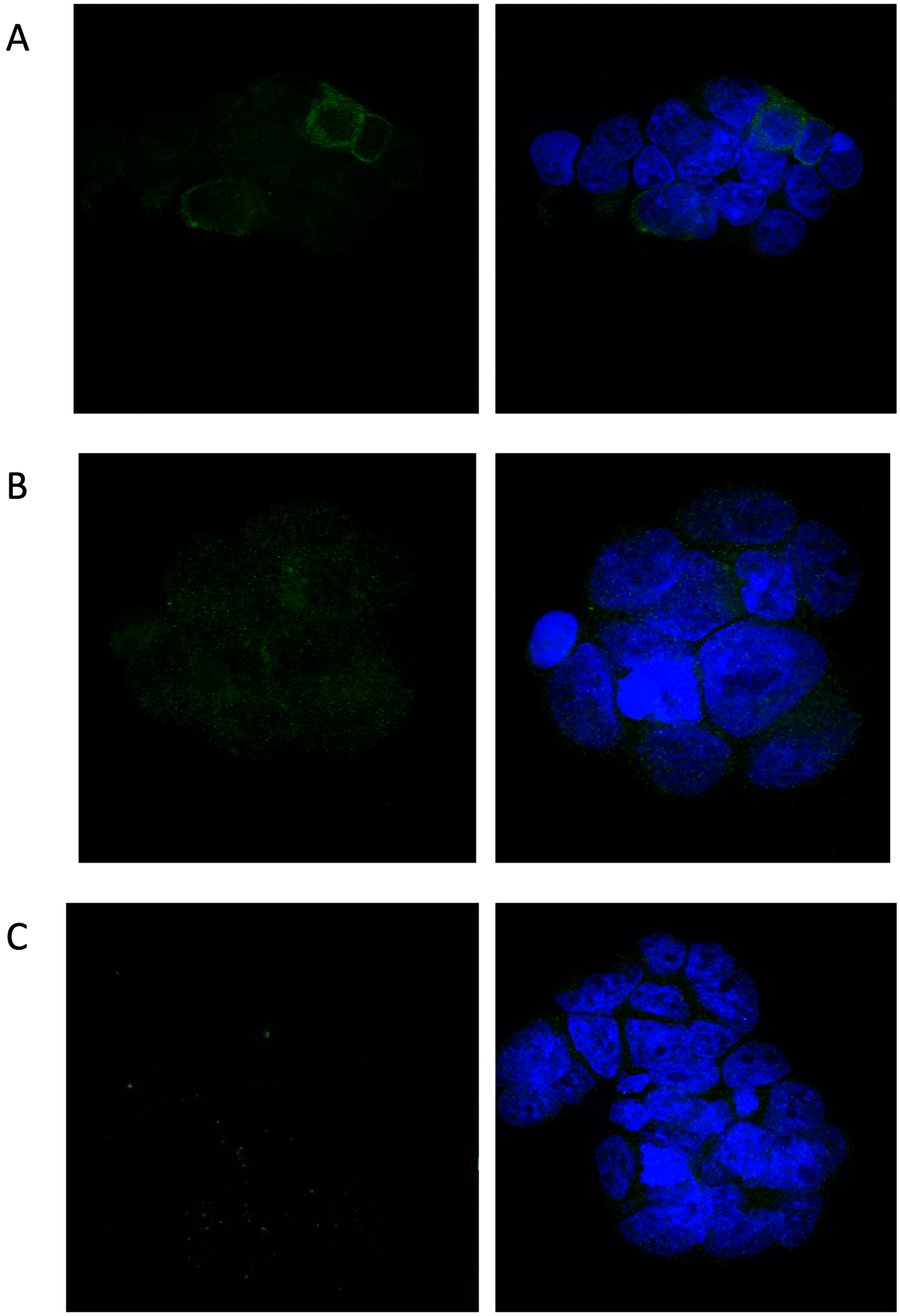


**Figure S2.** HEK293T cells transfected with pBudORF1-CH (A), pBudORF2-CH (B) (Addgene), and non-transfected control (C). Left panel: The immunofluorescence signal of ORF1p/ORF2p (green) in HEK293T cells. Right panel: The immunofluorescence signal of ORF1p/ORF2p (green) combined with the nuclei signal (DAPI, blue). Primary/secondary antibodies ORF1p: Anti-LINE-1 ORF1 (Abcam) + Goat anti Rabbit IgG AF488 (Abcam), ORF2p: Anti-LINE-1 ORF2 (Rockland) + Goat anti Chicken IgY AF488 (Invitrogen). Magnification: 40x.

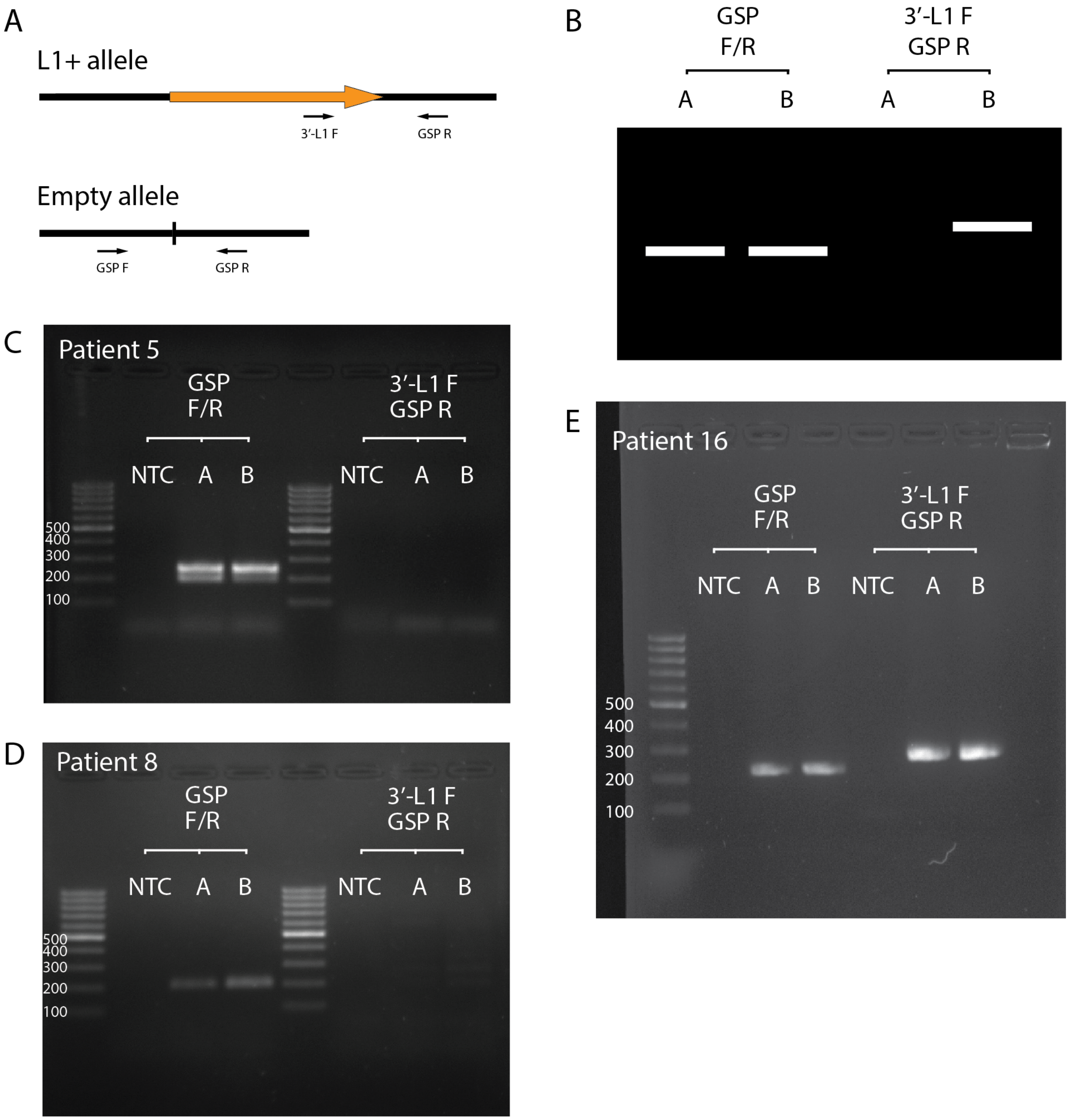


**Figure S3.** PCR validation of candidate L1 insertions detected by NGS. The approach is based on locus-specific PCR with primers designed for every candidate insertion. (A) We used a universal forward primer (3′-L1 F) specific for the 3′-end of every L1HS insertion (orange arrow) in combination with the genomic locus-specific reverse primer (GSP R) to capture the allele carrying novel insertion. In parallel, genomic locus-specific forward and reverse primers (GSP F and GSP R) were used to amplify an empty allele. Both PCR reactions were carried out for all consecutive samples of a patient or cell-line culturing condition. PCR products were then analyzed using agarose gel electrophoresis to reveal novel L1 insertions; a hypothetic positive result is illustrated (B). The results for the three MDS patients (5, 8, 16) with candidate insertions in the samples taken during Aza therapy turned out negative (C-E). Supplementary Tables S4 and S5 list PCR primers for all validation reactions. NTC: non-template control; A: before Aza; B: during Aza therapy.
