## Supplementary Methods for "Mobile retroelements induced by hypomethylating agents are restricted to transpose in myeloid malignancies"

**Fluorescence Microscopy**

**Culture Slide Preparation**

The sterilized coverslips with a diameter of 13 mm and a thickness of 0.17 mm (9161065, Menzel) were carefully inserted into the wells of the 24-well plate (92024, TPP). Next, the coverslips in the wells were coated with poly-L-lysine 0.01% (P4832, Sigma-Aldrich) and left to dry in a laminar box. The coverslips in the wells were seeded with the cells of the targeted cell line at a volume of 200 µl per well.

**Intracellular staining**

After completion of the cell line culture, the medium was removed from the wells and the cells were washed PBS and fixed with 4% Formaldehyde (F8775, Sigma-Aldrich) at RT for 10 min. Then, washing cells 3 times with PBS was followed by blocking with 3% IgG-free BSA blocking buffer (001-000-161, Jackson ImmunoResearch) with the addition of 0.25% Triton X-100 (SIALX100, Sigma-Aldrich) at RT for 1 hour. The cells were incubated in the corresponding primary antibody at RT for 1 hour, washed with 1× PBS and further incubated in the secondary antibody in dark at RT for ½ hour. For FL microscopy of both transfected and treated cells, the following antibodies were used: Anti-LINE-1 ORF1p (ab246320, Abcam) and Anti-LINE-1 ORF2 (200-901-B20S, Rockland) as primary antibodies, and Goat Anti Rabbit IgG AF488 (150077, Abcam) and Goat Anti Chicken IgY AF488 (A-11039, ThermoFisher) as secondary antibodies. After incubation, wells containing cells were washed 4 times with PBS followed by staining of nuclear DNA of cells with Hoechst stain (H1399, ThermoFisher) at RT for 2 min and washing once with PBS and once with distilled water. Finally, individual stained coverslips with cells were carefully transferred with tweezers to slides with prepared mounting medium (S3023, DAKO). The slides were detected the day after the mounting medium dried.

**Detection using a confocal microscope**

Fluorescence detection of ORF1p and ORF2p proteins was performed using a ZEISS 700LSM confocal microscope with plan-apochromat 40x oil objective lens. Individual images were detected using two filters with a wavelength of 405 and 488 nm and 1 AU pinhole and assessed in ZEN 2009 software.

**Preparation of ORF1 and ORF2 Plasmids and transient transfection**

*E.coli* DH5alpha with pBudORF1 (Addgene) and pBudORF2 (Addgene) were transferred on LB plate with Bleocin 1 mg/ml according to the manufacturer's protocol and cultured overnight at 37 ºC. Individual positive colonies were inoculated into individual Erlenmeyer flasks with

200 ml of culture LB medium and 1 mg/ml Bleocin using sterile loops. Culturing in flasks was carried out on a shaker at 300 rpm and 37 ºC until the next day. The individual bacterial suspensions were then centrifuged at 5000 g and 4 ºC for 15 minutes. The obtained bacterial pellets were processed and isolated using EndoFree Plasmid Maxi Kit (Qiagen) according to manufacturer's protocol. The resulting concentration and purity of the plasmid DNA was measured using a spectrophotometer at 260 and 280 nm.

**Transient transfection for FL microscop**y was performed directly on coverslips coated with poly-L-lysine (see above) placed in 24-well plates using the transfection reagent Polyethylenimine (PEI; Polysciences) according to the manufacturer's protocol. Briefly, the day before transfection, HEK293T cells were seeded in individual wells of a 24-well plate with coverslips at a cell density of $0.5 \times 10^6$/ml with 1 ml of DMEM/F12 culture medium (PAN-biotech) + 10% ultra-low IgG FBS (PAN-biotech). On the day of transfection, the culture medium was replaced with DMEM/F12:H2O transfection medium in a 1:1 ratio with the addition of 300 µl serum-free 2mM Glutamine (PAN-biotech). The premix was added to the individual wells with prepared transfection media after being left in the box at RT for half an hour at a DNA:PEI ratio (3:9 ųg) of 150 ųl. The transfected cells were incubated at 37 ºC and 5% $CO_2$. After 4 hours, the transfection medium was replaced with fresh 1ml DMEM/F12 medium + 10% ultra-low IgG FBS and plates were incubated for additional 72 hours. FL staining of the slide was carried out after PBS wash as described above.

**Transient transfection for the detection of ORF1p and ORF2p proteins by flow cytometry and western blots** was performed in 6-well plates (TPP) without the use of coverslips. The transfection procedure in the 6-well plates was identical to the transfection procedure in the 24-well plates. Given the larger well area of the 6-well plates compared to the 24-well plates, the volumes were increased accordingly, and 3 ml of HEK293T cells were used, the volumes of culture medium and transfection medium were 1 ml and 500 µl, respectively. After three days of culturing in 3 ml of fresh medium, each well was carefully washed with PBS. Finally, cells were then washed out with PBS supplemented with 15mM EDTA and transferred into 1.5 ml Eppendorf tubes.

**Library preparation for Next generation sequencing**

For the L1 flanks library preparation, 30 ng of gDNA was digested in 10 µL of 1× FD buffer with 5U of FspBI and 5U of TaqI (all Thermo Fisher Scientific, Waltham, MA, USA) for 30 min at 37 °C. For adapter ligation, fragmented DNA was diluted in 20 µL of 1× FD Buffer with 20 µmol of ATP (Thermo Fisher Scientific), 50 pmol of stem-loop (SL) adapters (St19N10hook_v2 and St19N10hook_TaqI) (see Table 1 for oligonucleotide sequences), 10U of T4 DNA ligase (Thermo Fisher Scientific), and incubated at 22 °C for 30 min. Next, 50 pmol of anti-adapters (antiHook-TA and antiHook-TaqI), additional 5U of FspBI and TaqI endonucleases were added, and the mixture was incubated at 22 °C for 30 min and 37 °C for additional 30 min. AntiSL-adapters inactivate SL-adapters, and endonucleases decrease the number of ligated chimeric molecules. The ligation reaction product was purified with 0.8 V of Ampure XP beads (Beckman Coulter, Brea, CA, USA), and used in the 1st PCR reactions containing 1× Encyclo Buffer, 1× Encyclo polymerase, 200 µM of each dNTP (all Evrogen, Moscow, Russia), 0.2 µM of L1HS-specific primer (St19roko-L1HS), and primer St19okor corresponding to the part of SL-adapter before the UMI. The amplification profile was: 95 °C for 2 min, followed by 12 cycles of 20 s at 95 °C, 20 s at 65 °C, and 1 min at 72 °C with a ramp rate of 1 °C/s. A total of 2 µL of the obtained 1st PCR product was added to the 2nd PCR reactions containing 1× Encyclo polymerase in 1× Encyclo Buffer, 200 µM of each of dNTP, 0.2 µM of each of the second PCR primers (KorNxt-L1HS – L1HS specific and korNxtSt19okor – adapter-specific). The amplification profile was: 2 min at 95 °C followed by 10 cycles of 20 s at 95 °C, 20 s at 60 °C and 1 min at 72 °C. A total of 2 µL of the 2nd PCR product was used in the third indexing PCR 25 µL reaction containing 200 µM of each dNTP, 1µl of Illumina Unique Dual indexing primers mix, and 1× Encyclo polymerase in 1× Encyclo Buffer amplified for 12 cycles using the same amplification profile as for the 2nd PCR. The indexing PCR products were purified with Ampure XP beads and mixed equimolarly for sequencing. Libraries were sequenced on Illumina NextSeq 550 (Illumina Inc, San Diego, CA, USA), paired-end 150 + 150.

**Table 1.** Oligonucleotides used for library preparation.

| Name | Sequence | Description |
|---|---|---|
| **Adapter ligation** | | |
| St19N10hook_v2 | TAAGCACGACGCCTGCGTGCTGCGGNNNNNNNNNNAGGCGTCGTGCT | SL-Adapter |
| St19N10hook_TaqI | CGTGCACGACGCCTGCGTGCTGCGGNNNNNNNNNNAGGCGTCGTGCA | SL-Adapter |
| antiHook-TA | pTAGTGTGCTCGTAGTCAAAAGACTACGAGCACAC | AntiSL-adapter |
| antiHook-TaqI | pCGGTGTGCTCGTAGTCAAAAGACTACGAGCACAC | AntiSL-adapter |
| **1st PCR** | | |
| St19okor | GCGTGCTGCGG | Adapter-specific primer |
| St19roko-L1HS | CGTCGTGCGTAGATGACAC | L1HS-specific primer |
| **2nd PCR** | | |
| korNxtSt19ok | TCGGCAGCGTCAGATGTGTATAAGAGACAGCGTGCTGCGG | Adapter-specific primer |
| KorNxt-L1HS | CGTGGGCTCGGAGATGTGTATAAGAGACAGCATGTACCCTAAAACTTAGA | L1HS-specific primer |
