## Supplementary Tables 1-5 for "Mobile retroelements induced by hypomethylating agents are restricted to transpose in myeloid malignancies"

**Table S1**. Cell lines used for experiments. All tumor cell lines were screened for ORF1p expression in the initial Western blot analysis. Flow cytometry and fluorescence microscopy were carried out on DAMI and HL-60 myeloid cell lines, with MCF-7 used as a positive control. HEK293T/17 cells were used for ORF1/2p overexpression.

| Cell line | Cell type origin | Origin | Catalogue No. | Cultivation medium | Method |
|---|---|---|---|---|---|
| MCF-7 | breast adenocarcinoma | DSMZ | ACC 115 | RPMI/DMEM* | initial WB screen, WB, FC, IF |
| SW-48 TP53 -/- | colorectal adenocarcinoma | Sigma-Aldrich | CLLS1007 | DMEM | initial WB screen |
| SW-48 | colorectal adenocarcinoma | ATCC | CCL-231 | DMEM | initial WB screen |
| H1299 | non-small cell lung carcinoma | ATCC | CRL-5803 | RPMI | initial WB screen |
| DAMI | treatment-related erythroleukemia after Hodgkin lymphoma | ATCC | CRL-9792 | RPMI/DMEM* | initial WB screen, WB, FC, IF |
| HL-60 | acute myeloid leukemia | DSMZ | ACC 3 | RPMI/DMEM* | initial WB screen, WB, FC, IF |
| MOLM-13 | secondary acute myeloid leukemia relapsed from myelodysplastic syndrome | DSMZ | ACC 554 | RPMI | initial WB screen |
| BV-173 | B cell precursor leukemia derived from blast crisis of chronic myeloid leukemia | DSMZ | ACC 20 | RPMI | initial WB screen |
| NALM-16 | B-cell precursor acute lymphoblastic leukemia | DSMZ | ACC 680 | RPMI | initial WB screen |
| NALM-6 | B-cell precursor acute lymphoblastic leukemia | DSMZ | ACC 128 | RPMI | initial WB screen |
| SU-DHL-4 | diffuse large B-cell lymphoma | DSMZ | ACC 495 | RPMI | initial WB screen |
| MINO | mantle cell lymphoma | DSMZ | ACC 687 | RPMI | initial WB screen |
| MAVER-1 | mantle cell lymphoma | DSMZ | ACC 717 | RPMI | initial WB screen |
| WSU-NHL | B-cell lymphoma, nodular histiocytic | DSMZ | ACC 58 | RPMI | initial WB screen |
| RAJI | Burkitt lymphoma | DSMZ | ACC 319 | RPMI | initial WB screen |
| HEK293T/17 | human embryonic kidney, contains the SV40 T-antigen | ATCC | CRL-11268 | DMEM | WB, FC, IF |

*WB: Western blot; FC: flow cytometry; IF: fluorescence microscopy*

** Cultivation in DMEM was used for experiments involving fluorescent microscopy or flow cytometry.*

**Table S2**. A cohort of MDS patients tested for L1 *de novo* insertions gained during Aza treatment.

| Patient ID | Sample order | Sampling date | Time from previous sampling (mo) | Material | Number of Aza cycles at sampling | Total number of Aza cycles | Azacytidine treatment - detail | TP53 status (mutations >1% VAF) |
|---|---|---|---|---|---|---|---|---|
| 1 | A | 10.10.2012 | - | BM | 0 | 22 | before Aza | wt |
| 1 | B | 09.09.2015 | 35 | BM | 22 | 22 | sAML | wt |
| 2 | A | 03.01.2018 | - | BM | 0 | 16 | before Aza | c.721T>G p.Ser241Ala 54.7% |
| 2 | B | 06.02.2019 | 13 | BM | 14 | 16 | during Aza | c.721T>G p.Ser241Ala 36.5% |
| 3 | A | 15.05.2019 | - | BM | 0 | 14 | before Aza | c.832C>T p.Pro278Ser 69.2% |
| 3 | B | 13.05.2020 | 12 | BM | 11 | 14 | during Aza | c.832C>T p.Pro278Ser 31.7% |
| 4 | A | 11.03.2020 | - | BM | 0 | 34 | before Aza | wt |
| 4 | B | 12.05.2021 | 14 | BM | 12 | 34 | during Aza | wt |
| 5 | A | 21.01.2020 | - | BM | 0 | 9 | before Aza | not done |
| 5 | B | 04.05.2020 | 3 | BM | 3 | 9 | during Aza | wt |
| 5 | C | 09.02.2021 | 9 | BM | 6 | 9 | relapse after alloTx, sAML 1/2021 | wt |
| 6 | A | 26.03.2014 | - | BM | 0 | 10 | before Aza | wt |
| 6 | B | 10.12.2015 | 20 | BM | 6 | 10 | during Aza | wt |
| 7 | A | 05.12.2019 | - | BM | 0 | 5 | before Aza | wt |
| 7 | B | 30.04.2020 | 5 | BM | 4 | 5 | during Aza | wt |
| 8 | A | 15.01.2014 | - | BM | 0 | 5 | before Aza | wt |
| 8 | B | 20.06.2014 | 5 | BM | 5 | 5 | during Aza, sAML 7/2014 | wt |
| 9 | A | 08.06.2017 | - | BM | 0 | 14 | before Aza | wt |
| 9 | B | 02.05.2018 | 11 | BM | 9 | 14 | during Aza, sAML 10/2018 | wt |
| 10 | A | 14.03.2018 | - | BM | 0 | 14 | before Aza | wt |
| 10 | B | 06.01.2021 | 34 | BM | 10 | 14 | during Aza, sAML 4/2020 | wt |
| 11 | A | 28.02.2018 | - | BM | 0 | 10 | before Aza | wt |
| 11 | B | 13.02.2019 | 11 | BM | 10 | 10 | during Aza | wt |
| 12 | A | 08.04.2015 | - | BM | 0 | 20 | before Aza | p.Val216Met 1.4% |
| 12 | B | 31.01.2018 | 34 | BM | 19 | 20 | during Aza | p.Val216Met 3.6% |
| 12 | C | 15.05.2019 | 15 | BM | 20 | 20 | sAML, after Aza and lenalidomide | p.Val216Met 21.2% |
| 13 | A | 28.03.2018 | - | BM | 0 | 14 | before Aza | c.733G>A p.Gly245Ser 18.6%; c.707A>G p.Tyr236Cys 2.5% |
| 13 | B | 28.08.2019 | 17 | BM | 14 | 14 | relapse (progression of anemia) | c.733G>A p.Gly245Ser6.5%; c.707A>G p.Tyr236Cys 4.5% |
| 13 | C | 24.11.2020 | 15 | BM | 14 | 14 | after Ara-C | c.733G>A p.Gly245Ser 4.0% |
| 14 | A | 03.03.2016 | - | BM | 0 | 7 | before Aza | wt |
| 14 | B | 20.07.2016 | 5 | BM | 4 | 7 | sAML | wt |
| 15 | A | 28.12.2012 | - | BM | 0 | 14 | before Aza | wt |
| 15 | B | 04.06.2014 | 17 | BM | 14 | 14 | after Aza (sAML 9/2014) | wt |
| 16 | A | 01.03.2012 | - | BM | 0 | 53 | before Aza | wt |
| 16 | B | 31.05.2017 | 63 | BM | 46 | 53 | during Aza (sAML 5/2018) | wt |
| 17 | A | 10.07.2013 | - | BM | 0 | 13 | before Aza | c.637C>T p.Arg213Ter 34%; c.430C>T p.Gln144Ter 18% |
| 17 | B | 21.08.2014 | 13 | BM | 12 | 13 | relapse (increase in blast counts, progression of pancytopenia) | c.637C>T p.Arg213Ter 23%; c.430C>T p.Gln144Ter 11% |

*sAML: progression to acute myeloid leukemia*

**Table S3**. Antibodies used for Western blot analysis (WB), flow cytometry measurements (WB), and immunofluorescence staining for confocal microscopy (IF).

| Antibody | Manufacturer | Cat. no. | Type | Species | Application |
|---|---|---|---|---|---|
| Anti-LINE-1 ORF1p antibody [EPR22227-54] - BSA and Azide free | Abcam | ab246320 | monoclonal | rabbit | WB, FC, IF |
| L1/ORF2 Antibody | Rockland | 200-901-B20S | polyclonal | chicken | WB, FC, IF |
| ORF1p (D3W9O) Rabbit mAb | CellSignalling | 88701 | monoclonal | rabbit | WB |
| Anti-PCNA Antibody, clone PC10 | Millipore | MAB424R | monoclonal | mouse | WB |
| Anti-β-Actin antibody | Sigma-Aldrich | A5441 | monoclonal | mouse | WB |
| Anti-mouse IgG HRP | Cell Signaling | 7076 | polyclonal | horse | WB |
| Goat Anti-Rabbit IgG H&L (Alexa Fluor® 488) | Abcam | ab150077 | polyclonal | goat | FC, IF |
| Anti-Rabbit IgG HRP | Cell Signaling | 7074 | polyclonal | goat | WB |
| Goat anti-Chicken IgY (H+L) Secondary Antibody, Alexa Fluor™ 488 | Invitrogen | A-11039 | polyclonal | goat | FC, IF |
| Goat Anti-Chicken IgY H&L (HRP) | Abcam | ab6877 | polyclonal | goat | WB |

*WB: Western blot; FC: flow cytometry; IF: fluorescence microscopy*

**Table S4**. A list of candidate insertions detected using L1-targeted amplicon-based NGS protocol and validated using site-specific PCR in DAMI and HL-60 cell lines subjected to 28 days lasting Aza-dC treatment.

| Cell line | Replicate | Aza-dC concentration (uM) | Day of cultivation | Insertion genomic localization | Read count | UMI count | Forward flank primer | Reverse flank primer | Forward L1 primer | PCR result |
|---|---|---|---|---|---|---|---|---|---|---|
| DAMI | 1 | 0,5 | 14 | chr5: 110,920,230 | 15 | 6 | TTAGCAAATAGTTGGCCCTTTT | ATGGTTGCAAATTCAGCCCATA | TGGGAGATATACCTAATGCTAGATGACAC | negative |
| DAMI | 1 | 0,5 | 28 | chr2: 228,070,005 | 31 | 3 | CTGGCAGTTGTGACTTGCAT | TTTTACTGCACAGGGGTTGG | TGGGAGATATACCTAATGCTAGATGACAC | negative |
| DAMI | 1 | 0,5 | 7 | chr18: 48,544,088 | 11 | 2 | AAGCTTGGTCTGTGGCTTTC | AGTAGCCCCAGTGTCAACCA | TGGGAGATATACCTAATGCTAGATGACAC | negative |
| DAMI | 1 | 0 | 7 | chr4: 23,894,966 | 20 | 2 | TGTTCAACTCGGTTGTATTTGTG | ATGCCTTGTTCAGTTCCACTT | TGGGAGATATACCTAATGCTAGATGACAC | negative |
| DAMI | 1 | 0 | 7 | chr4: 177,188,221 | 19 | 3 | AGCAAATCCAGTCAGCAAGC | TGTTTTTGGTCAGGGGGCAT | TGGGAGATATACCTAATGCTAGATGACAC | negative |
| DAMI | 2 | 0 | 14 | chr17: 4,354,041 | 12 | 2 | GGTCTTCCACAAACGTCAACC | CCAGTCTGGTTCATGGAGGG | TGGGAGATATACCTAATGCTAGATGACAC | negative |
| DAMI | 2 | 0 | 3 | chr1: 7,703,623 | 21 | 3 | GCTTAAGGGCTCTGCAAATG | CTGACCCCAAGACCTGTGAT | TGGGAGATATACCTAATGCTAGATGACAC | negative |
| DAMI | 2 | 2 | 28 | chr4: 20,796,978 | 12 | 2 | CCGTGTTCAAAAACACAGCA | GGTGACACTTGAGCCCTTTC | TGGGAGATATACCTAATGCTAGATGACAC | negative |
| HL60 | 1 | 2 | 14 | chr6: 105,077,816 | 12 | 3 | TGGTAATTTGCCAGATATCATCC | GGCATGATACAAAGTGGCAAT | TGGGAGATATACCTAATGCTAGATGACAC | negative |
| HL60 | 1 | 2 | 7 | chr6: 132,078,310 | 13 | 3 | CCGGGTTAGATGCTGATGAT | TGCATAGGACTGAAAACATCCA | TGGGAGATATACCTAATGCTAGATGACAC | negative |
| HL60 | 2 | 0,5 | 3 | chr8: 24,223,576 | 32 | 5 | CCAAGGCCCCATGTTAGATA | GGGTGGGAAAACAAGGATTT | TGGGAGATATACCTAATGCTAGATGACAC | negative |
| HL60 | 2 | 0 | 3 | chr3: 105,036,561 | 13 | 2 | CCTGGTCCTCAAAAATTCCA | CTTGGCACCTGCCTACATTT | TGGGAGATATACCTAATGCTAGATGACAC | negative |
| HL60 | 2 | 2 | 14 | chr13: 96,139,793 | 29 | 6 | TTCTTCATGAGTTTTGCATAGGA | CCCAAGTACTGCACCCAAAT | TGGGAGATATACCTAATGCTAGATGACAC | negative |
| HL60 | 2 | 2 | 3 | chr19: 3,193,795 | 17 | 5 | TGCACACGAAAGACCAGAAC | CCCTTCTCCGTAAGGACACC | TGGGAGATATACCTAATGCTAGATGACAC | negative |

**Table S5**. A list of candidate insertions detected using L1-targeted amplicon-based NGS protocol and validated using site-specific PCR in MDS patient samples after Aza treatment.

| Patient ID | Sample with candidate de novo insertion | Compared against sample | Insertion genomic localization | Forward flank primer | Reverse flank primer | Forward L1 primer | PCR result |
|---|---|---|---|---|---|---|---|
| 5 | B | A | chr12: 77,122,549 | TAATAACTGAGGCGGCTGCT | CCTGTGGTTTCACTTTTTCCA | TGGGAGATATACCTAATGCTAGATGACAC | negative |
| 8 | B | A | chr9: 13,050,004 | TTGGAATACACCCTTCACGAG | TTGCAAATGAAATGCAGCAATA | TGGGAGATATACCTAATGCTAGATGACAC | negative |
| 16 | B | A | chr21: 32,361,376 | TGCCTTGAAACAAAAACTGC | TTTGCCAGACCTGAACACTG | TGGGAGATATACCTAATGCTAGATGACAC | negative |
